# Novel PSD95 reporter mice reveal medium spiny neuron subtype-specific synapse loss in PD and L-dopa induced dyskinesia and identify microglia mediated synapse removal as a therapeutic target for dyskinesia

**DOI:** 10.64898/2026.07.05.736610

**Authors:** Peggy Rentsch, Jessica Irving, Illya Conn, Kathryn J Laloli, Luke T Milham, Sandy Stayte, Bryce Vissel

**Author notes:** Correspondence Bryce Vissel.

## Abstract

**Background:** L-Dopa remains the primary treatment for Parkinson’s disease (PD), but chronic administration frequently leads to L-Dopa-induced dyskinesia (LID). While D1 and D2 medium spiny neuron (MSN) specific structural changes on the spine level have been observed in the striatum of PD and LID, studying microglia mediated synapse loss has not been done to date.

**Methods:** Here we generated novel reporter mice by crossing floxed PSD95c(mCherry/eGFP) mice with D1-Cre and D2-Cre lines, producing D1-PSD95-EGFP and D2-PSD95-EGFP strains for MSN-specific synapse visualization. Using the 6-OHDA mouse model of PD and LID we assessed microglia mediated MSN subtype specific synapse loss in these mice while PLX3397 was used to investigate effects of microglia depletion and repopulation on LID development and synapse loss.

**Results:** Both D1- and D2-MSNs exhibited significant PSD95 synapse loss in PD, with D1-MSN loss further exacerbated in LID. Microglia displayed increased phagocytic activity and accumulated PSD95 material within lysosomes, particularly in LID. PLX3397-mediated microglial depletion reduced LID severity and preserved D1-MSN synapses. A depletion and repopulation paradigm attenuated LID severity, preserved D1-MSN synapses, and reduced synaptic material within microglia.

**Conclusions:** Microglia-mediated synapse loss in MSN subtypes contributes to PD and LID pathogenesis. Pharmacological microglial depletion and repopulation mitigate synapse loss and dyskinesia, highlighting microglial turnover as a promising therapeutic strategy for LID.

## Introduction

Although L-Dopa remains the primary therapy for Parkinson’s disease (PD), chronic administration leads to abnormal involuntary movements (AIMs) known as L-Dopa induced dyskinesias (LIDs) in more than 70% of PD patients within 10 years of treatment^1,2^. Beyond the well-characterized dopaminergic deficits, accumulating evidence indicates that profound synaptic remodelling in the striatum of the basal ganglia circuitry accompanies both PD and LID development^3^. The striatum is primarily composed of GABAergic medium spiny neurons (MSNs) of two subtypes, which form two pathways with opposing outcomes that operate in a push - pull manner to regulate motor control^4^. While D1 receptor expressing MSNs (D1-MSNs) are part of the direct pathway and promote action selection, D2 receptor expressing MSNs (D2-MSNs) form part of the indirect pathway and are thought to suppress contextually inappropriate actions. Given these contrasting functional outcomes, it is crucial to study synaptic or structural changes in an MSN subtype specific manner, as global analyses may offset opposing effects.

Previous work using 6-OHDA mouse models has consistently shown that dopamine depleted parkinsonian animals exhibit reduced spine density in indirect pathway D2-MSNs, and this spine loss is restored following L-Dopa treatment^5–9^. In contrast, findings for the direct pathway are less consistent. While some studies report spine loss in striatal D1-MSNs in dopamine depleted parkinsonian animals^5,6,9^, others report spine loss only occurs after prolonged L-Dopa treatment rather than as a direct consequence of the 6-OHDA lesion^7,8,10^. These discrepancies may reflect methodological differences, including variations in 6-OHDA injection sites or lesion development timeframes^9,11^. Furthermore, these studies consistently utilize bacterial artificial chromosome (BAC) transgenic mice expressing fluorescent markers in D1- or D2-MSNs (BAC-*Drd1a* or BAC-*Drd2* mice), together with dye-filling and two-photon laser scanning microscopy (2PLSM) to quantify spine density. While widely adopted, this approach may not capture smaller spines due to resolution constraints^10^, introduce a bias toward filling morphologically intact neurons, is neuron rather than wider region specific, and provides only a structural approximation of connectivity under the assumption that each spine contains a functional postsynaptic density. It would therefore be of interest for the PD and LID field to study functional synapse loss across the striatum in an MSN subtype specific manner, however, to date, tools required for such analysis were missing.

Meanwhile, in other neurodegenerative disorders, including Alzheimer’s (AD)^12–14^ and Huntington’s disease (HD)^15^, synapse loss has been extensively characterized, and accumulating evidence implicates microglial dysfunction as a key driver of this process. Similarly to these conditions, microglial activation is a well-established feature of PD^16–18^ and is increasingly recognized as contributing to the pathophysiology of LID^19–23^. In line with this, we have demonstrated that striatal microglia adopt a reactive, phagocytic phenotype in PD, a response that is further exacerbated in parkinsonian animals subjected to repeated L-Dopa treatment^23^.

To investigate microglia-mediated synapse removal in an MSN subtype-specific manner in PD and LID in the current study, we generated novel reporter mice by crossing floxed PSD95c(mCherry/eGFP)^24^ mice with D1-Cre and D2-Cre lines, resulting in D1-PSD95-EGFP and D2-PSD95-EGFP strains, respectively. Using these mice, we observed that both D1- and D2-MSNs exhibit loss of PSD95-containing synapses in PD, with synapse loss in D1-MSNs further exacerbated in LID. This synapse loss coincided with lysosomal compartments of microglia containing markedly more synaptic material in LID, implicating functional relevance. To directly assess the role of microglia, we depleted approximately 90% of striatal microglia using the FDA-approved CSF1R inhibitor PLX3397 (Pexidartinib)^25–27^, which resulted in both reduced LID severity and prevention of synapse loss on D1-MSNs. We next implemented a depletion - repopulation paradigm, aiming to replace reactive microglia arising during lesion development with newly repopulated cells. Strikingly, mice harbouring repopulated microglia exhibited attenuated LID severity, reduced PSD95 synapse loss in D1-MSNs and decreased synaptic material within microglia. Collectively, these findings demonstrate that striatal synapse loss in both MSN subtypes is a critical contributor to LID pathogenesis and that preventing microglia-mediated synapse removal represents a promising therapeutic strategy.

## Methods

### Animals

The B6;129-Tg(Drd1-cre)120Mxu/Mmjax (MMRRC:037156-JAX) and B6.FVB(Cg)-Tg(Drd2-cre)ER44Gsat/Mmucd (MMRRC:032108-UCD) mouse lines were originally obtained from the Mutant Mouse Resource and Research Centers (MMRRC). The PSD95^c(mCherry/eGFP)^ construct was generously provided by Professor Seth Grant (University of Edinburgh). Drd1-cre and Drd2-cre lines were subsequently crossed with floxed PSD95^c(mCherry/eGFP)^ mice to generate D1-PSD95-EGFP and D2-PSD95-EGFP lines, respectively, and all genetically modified mice used in experiments were homozygous for the floxed gene and heterozygous for the Cre gene. In these lines, endogenous PSD95 is tagged with eGFP in either D1 or D2 MSNs, enabling MSN subtype-specific synapse visualization. C57BL/6J controls, D1-PSD95-EGFP and D2-PSD95-EGFP mice of all sexes were bred and maintained at Australian BioResources (Moss Vale, Australia) until experimentation. During experiments, mice were kept in groups of up to five per cage and maintained on a 12-hour light/dark cycle with access to food and water *ad libitum*. Animal experiments were performed with the approval of the Garvan Institute and St. Vincent’s Hospital Animal Ethics Committee under approval numbers 18/37, 20/10 and 23/13 and in accordance with the Australian National Health and Medical Research Council animal experimentation guidelines and the Australian Code of Practice for the Care and Use of Animals for Scientific Purposes. All surgeries were performed under ketamine (8.7 mg/mL; Mavlab) and xylazil (2 mg/mL; Troy Laboratories) anaesthesia. The experimenter was always blinded to group and outcome assignment, and tissue collection and processing was performed in appropriate blocks.

### Primary Striatal Neuron Culture

Primary striatal neurons were isolated from E15-E17 embryos obtained from D1-PSD95-EGFP and D2-PSD95-EGFP mouse lines. Pregnant dams were euthanised by CO_2_ inhalation followed by cervical dislocation. Embryos were rapidly extracted and individual brains were collected in ice-cold harvesting solution (HBSS without Ca2^+^/Mg2^+^ [Gibco, Cat# 14175095]). For each embryo, cortical tissue was collected and snap-frozen for subsequent genotyping. The striatum was identified and bilaterally dissected under a dissecting microscope with meninges removed where feasible. Individual embryonic striatal tissue was processed separately to maintain identification throughout culture establishment.

Striatal tissue was enzymatically dissociated in harvesting solution containing 0.25% trypsin (Merck, Cat# T4799-5G) for 30 minutes at 37°C with 5% CO_2_. Trypsin was inactivated with harvesting solution supplemented with 10% foetal calf serum (Gibco, Cat# A5669701). Tissue was mechanically dissociated by trituration through a 200 μL pipette tip. Cell suspensions were centrifuged at 300 x g for 5 minutes and pellets were resuspended in plating solution (Neurobasal Plus Medium [Gibco, Cat# A3582901], 5% FCS, 1X B27 Plus [Gibco, Cat# A3653401], 1X HEPES [Gibco, Cat# 15630080]). Viable cell concentration was determined by trypan blue exclusion. Neurons from each embryo were plated separately at 10-15 x 10^4^ cells/mm² on individual glass coverslips that had previously been coated overnight with 0.05 mg/mL poly-D-lysine (Gibco, Cat# A3890401) at 37°C before aspiration. Three hours post-plating, an equal volume of plating medium was added to each well. At DIV1, the day after plating, 1 μM cytosine arabinoside (Thermo Fisher Scientific, Cat# 449560050) was added to inhibit glial proliferation. Equal volumes of maintenance medium (Neurobasal Plus Medium, 1X B27 Plus, 1X HEPES) were added to cultures at DIV4, DIV8 and DIV12.

### Genotyping

Total RNA was extracted from snap-frozen cortical tissue using a silica column RNA extraction kit (Invitrogen, Cat# 12183025) as per manufacturer’s instructions. Complementary DNA was synthesised using reverse transcription (Invitrogen, Cat# 18091050). Cre recombinase expression was determined by quantitative PCR using Cre-specific primers (Table 1) and interleukin-2 as an internal control. Thermocycling conditions were: 95°C for 10 minutes, followed by 40 cycles of 95°C for 15 seconds and 60°C for 60 seconds. Only cultures derived from Cre-positive embryos were retained for subsequent analysis.

**Table 1:**
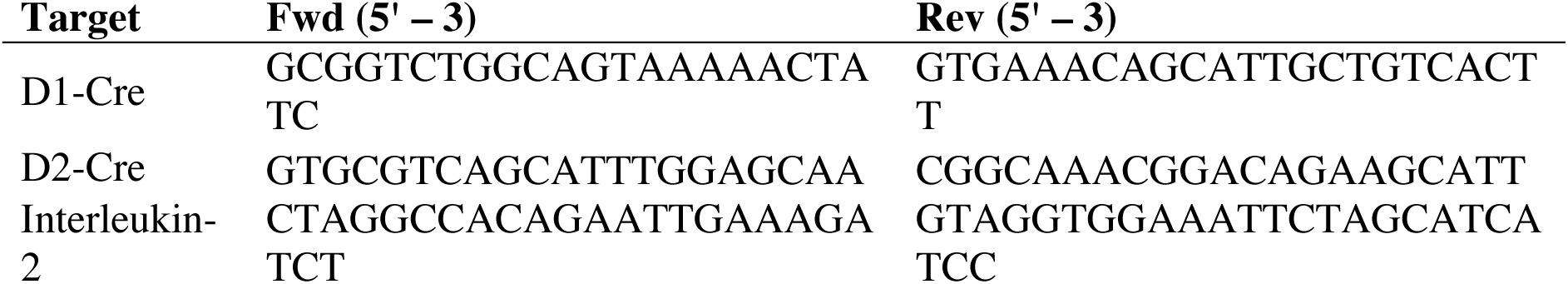
Primer nucleotide sequences for Cre genotyping.

### Immunocytochemistry

At DIV14, neurons were fixed with 4% paraformaldehyde for 15 minutes at room temperature. Cultures were permeabilised with 0.1% Triton X-100 (Thermo Fisher Scientific, Cat# A16046-AE) in phosphate-buffered saline for 30 min and blocked with 10% normal horse serum (Gibco, Cat# 16050130) for 1 hour at room temperature. Primary antibodies against dopamine receptor D1 (Merck, Cat# D2944, dilution 1/200), dopamine receptor D2 (Merck, Cat# AB5084P, dilution 1/200), and GFP (Abcam, Cat# ab13970, dilution 1/500) were applied in blocking solution overnight at 4°C. Following three washes in PBS, cultures were incubated with species-appropriate Alexa Fluor secondary antibodies (Invitrogen, dilution 1/500) for 1 hour at room temperature. Coverslips were mounted on glass slides using glycerol and imaged by using Leica Thunder widefield microscope equipped with a 63× Plan 1.4 NA oil immersion objective.

### Unilateral medial forebrain bundle (MFB) lesion

Anesthetized 8 week old mice were stereotaxically injected with 6-hydroxydopamine hydrobromide (0.2 μL of 15 mg/mL base-free 6-OHDA in 0.02% ascorbic acid; Sigma) into the right MFB (AP -1.2, ML -1.1, DV -5.3, relative to bregma), as previously described^28,29^.

### Abnormal involuntary movements (AIMs)

To induce LID, mice received repeated intraperitoneal injections of L-Dopa methyl ester (6 mg/kg; Sigma) and Benserazide-HCl (12.5 mg/kg; Sigma) in saline over a three-week period. To assess dyskinetic behaviour the AIMs scale described previously was used^28,30,31^. Axial, limb, and orolingual AIMs were scored on a 1–4 scale based on severity (amplitude) and duration (basic scale). A global AIM score was calculated by multiplying the basic and amplitude scores.

### Microglia depletion and repopulation

Mice received a chow diet (SF14-156, Speciality Feeds) containing 600 mg/kg PLX3397 (SelleckChem) *ad libitum* for 7 days to deplete microglia and in experiments assessing the anti-dyskinetic efficacy of PLX3397 mice remained on that for another 14 days until the end of the experiment. Control chow with the same base formula without PLX3397 was given to the control group. For repopulation experiments, mice were returned to the standard chow diet after 7 days and for the remainder of the experiments.

### Immunohistochemistry

The day after the last AIMs scoring session, mice were transcardially perfused with 4% paraformaldehyde 1 hour after L-Dopa administration. Brains were harvested, postfixed and processed as described previously^32^. For PSD95 synaptic puncta quantification 14 μm sections were slide mounted onto SuperFrost-plus slides (Menzel-Glaser) and coverslipped with Vectashield Hardset antifade mounting media with DAPI (Vector Laboratories). For microglia analysis 40 μm coronal brain sections were blocked with 3% BSA + 0.25% Triton-X-100 and then incubated in primary antibodies for 72 hours at 4 °C (Table 2). All sections were incubated in their respective secondary antibodies overnight at 4 °C followed by a counterstain with 4′,6-diamidino-2-phenylindole (DAPI; Life Technologies). Finally, sections were mounted onto SuperFrost-plus slides (Menzel-Glaser) and coverslipped with 50% glycerol mounting medium.

**Table 2:** Antibodies used for immunohistochemistry.

| **Antibody** | **Host Species** | **Concentration** | **RRID** |
| --- | --- | --- | --- |
| IBA1 | Rabbit | 1:1000 | RRID:AB_839504 |
| CD68 | Rat | 1:250 | RRID:AB_322219 |
| TH | Mouse | 1:1000 | RRID:AB_477569 |
| Anti-Rat 488 | Donkey | 1:500 | RRID:AB_2535794 |
| Anti-Mouse 555 | Donkey | 1:500 | RRID:AB_2536180 |
| Anti-Rabbit 647 | Donkey | 1:500 | RRID:AB_2536183 |

### PSD95 puncta analysis

Image stacks spanning 4-6 μm were acquired using either a DeltaVision OMX SR microscope equipped with a 60× Plan 1.42 NA oil immersion objective, with structured illumination reconstructions performed in Softworx software, or a Leica Thunder widefield microscope equipped with a 63× Plan-Apochromat 1.4 NA oil immersion objective, with Large Volume Computational Clearing applied using LAS X v3.8 software. Synaptic puncta were quantified in FIJI (ImageJ) using the SynQuant plugin. The number of synaptic puncta was then divided by total volume of the image stack. For each brain, 3–4 image stacks were acquired, and their mean value was considered as one biological replicate (n = 1).

### Microglia engulfment analysis

Image stacks spanning 10 μm were acquired using a Leica Thunder widefield microscope equipped with a 63× Plan-Apochromat 1.4 NA oil immersion objective. Large Volume Computational Clearing was applied during acquisition using LAS X v3.8 software. Image analysis was performed in ImageJ. Only microglia for which the entire cell was captured within the image stack were included for analysis. For subsequent processing, images were cropped to isolate individual microglia. Image stacks were converted to maximum intensity projections. Each fluorescence channel was subjected to automatic thresholding using the Moments (dark) algorithm and converted to a binary mask. For the IBA1 and CD68 channels, the total area of positive particles was quantified, and CD68 area was expressed as a percentage of the IBA1-positive area. To quantify PSD95 material within lysosomes, a colocalization mask was generated by computing the overlap between PSD95 and CD68 channels. The total area of positive particles within this mask was measured, and PSD95 was expressed as a percentage of the total CD68-positive area.

### Stereology

Microglial counts within the entire striatum and TH-positive neurons in the substantia nigra pars compacta (SNpc) were estimated using stereological analysis with the optical fractionator method (Table 3) implemented in Stereo Investigator software, version 7 (MBF Bioscience). Sampling precision was assessed by calculating the coefficient of error according to the approach described by Gundersen and Jensen^33^. Errors ≤0.10 were acceptable. Data are presented as number of cells per striatal volume (both measures determined using Stereo Investigator) for a more accurate assessment, as striatal volume differs between subjects dependent on the start (1 in 12) of the serial section.

**Table 3:** Stereological parameters.

| **Marker** | **No. of sections counted** | **Section Interval** | **Bregma coordinates** | **Guard Zone** | **Dissector Height** | **Counting Frame Size** | **Grid Size** |
| --- | --- | --- | --- | --- | --- | --- | --- |
| IBA1/CD68 | 7 | 12 | 1.6mm to -1.6mm | 5$\mu\text{m}$ | 10$\mu\text{m}$ | 100$\mu\text{m}$ x 100$\mu\text{m}$ | 333$\mu\text{m}$ x 333$\mu\text{m}$ |
| TH | 10 | 3 | -3.8mm to -2.6mm | 5$\mu\text{m}$ | 10$\mu\text{m}$ | 150$\mu\text{m}$ x 150$\mu\text{m}$ | 190$\mu\text{m}$ x 190$\mu\text{m}$ |

### Statistics

All statistical analyses were performed using Prism 6 (GraphPad). Shapiro-Wilk tests were performed on all data sets to assess normality, before analysing data either with parametric or non-parametric tests. For normally distributed data, differences between means were assessed, as appropriate, by t-tests, one- or two-way ANOVA with or without repeated measures, followed by Bonferroni post hoc analysis and not normally distributed data were assessed by Mann-Whitney test. All data is presented as mean ± standard error of the mean (SEM). For all statistical tests, *p*≤0.05 was assumed to be significant.

## Results

### Generation and Validation of MSN Subtype-Specific PSD95-EGFP Reporter Mice

As MSNs and their synaptic connections are intermingled within the striatum, we generated novel reporter mouse lines to study MSN subtype-specific synapse loss in PD and LID. Upon Cre-recombination in these mice, PSD95 is tagged with EGFP selectively in either D1-expressing MSNs (D1-PSD95-EGFP; Figure 1A) or D2-expressing MSNs (D2-PSD95-EGFP; Figure 1B). To validate MSN subtype specificity, we cultured neurons from D1-PSD95-EGFP mice and observed that EGFP signal colocalized exclusively with D1+ neurons, with PSD95-EGFP puncta distributed along dendritic spines (Fig. 1C). Similarly, cultures derived from D2-PSD95-EGFP mice exhibited PSD95-EGFP puncta restricted to D2+ neurons and localized along dendrites (Fig. 1D). These reporter lines provide a platform for quantifying MSN-specific synapse loss under pathological conditions.

**Figure 1:**
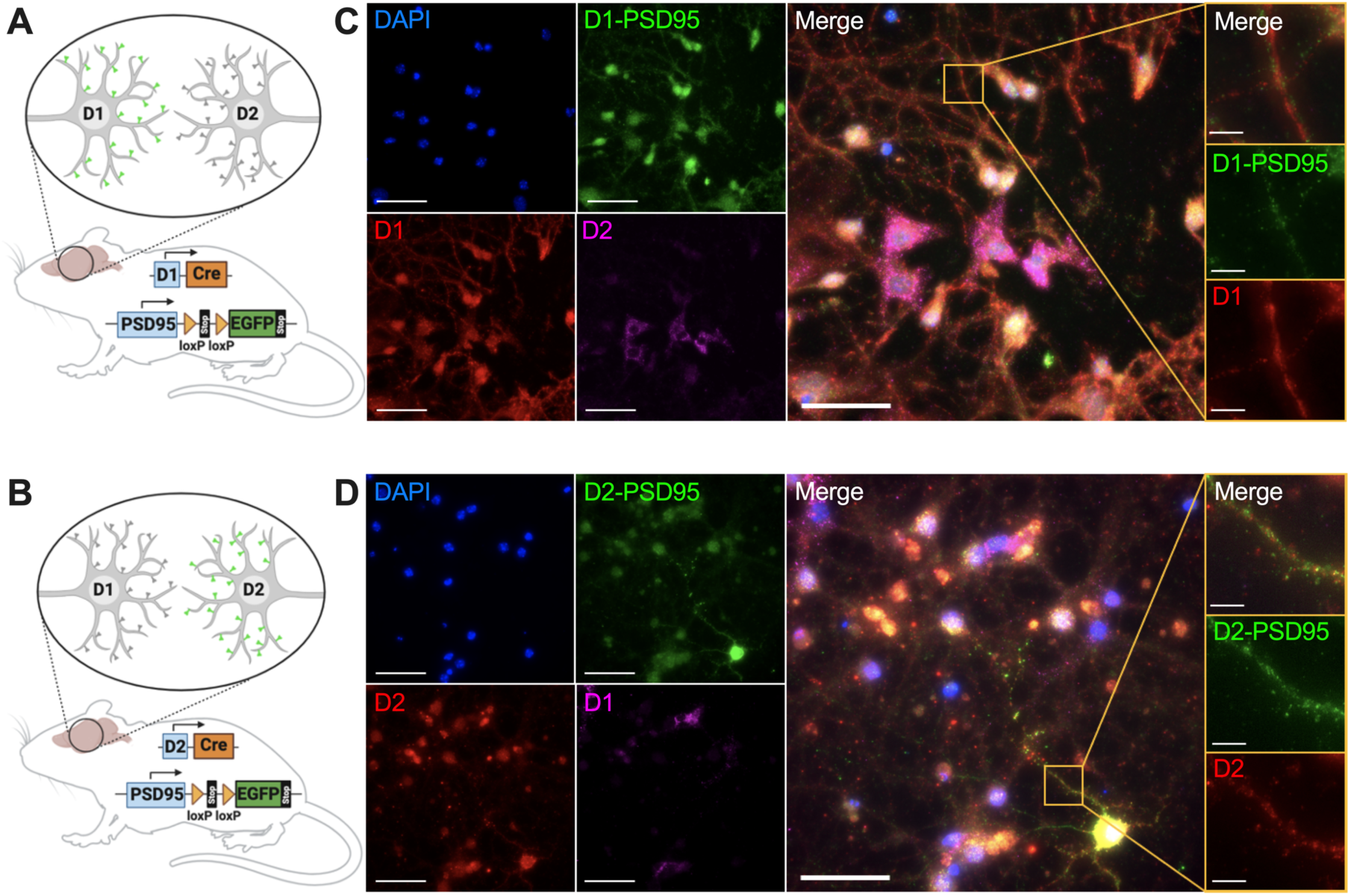
Generation and validation of MSN subtype-specific PSD95-EGFP reporter mice. Genetic design of Cre-loxP strategy to generate (A) D1-PSD95-EGFP and (B) D2-PSD95-EGFP fluorescent reporter lines to selectively tag PSD95 in D1- or D2-expressing MSNs. (C) Fluorescent images of cultured striatal neurons from D1-PSD95-EGFP mice demonstrated colocalization of PSD95-EGFP puncta with D1 immunostaining, but not D2 staining. High-magnification views revealed PSD95-EGFP puncta distributed along D1+ dendritic spines. (D) Fluorescent images of cultured striatal neurons from D2-PSD95-EGFP mice showed colocalization of PSD95-EGFP puncta with D2 immunostaining, but not D1 staining. Zoomed-in images confirmed PSD95-EGFP puncta localized along D2+ dendrites. Scale bar = 50μm and scale bar = 5μm for zoomed in images.

### PD-induced PSD95 synapse loss on D1-MSNs is exacerbated in LID and is associated with increased microglial lysosomal PSD95 content

To assess disease stage-dependent synapse loss, we collected brain tissue from mice with distinct phenotypes. Mice underwent ascorbic acid (naïve group) or 6-OHDA lesion surgery. Three weeks post-lesion, one subset was classified as PD (6-OHDA, no L-Dopa), and the others received repeated L-Dopa/Benserazide injections for three weeks. Based on global AIMs scores (Supplementary Fig. 1A), only mice scoring >40 were categorized as dyskinetic and included for analysis. Tissue was collected at study completion, and extensive DA neuron loss was confirmed in PD and LID groups (Supplementary Fig. 1B).

In the ipsilateral striatum of D1-PSD95-EGFP mice, parkinsonian animals exhibited a statistically significant reduction in PSD95-EGFP puncta compared to naïve controls (p < 0.01), and this loss was further exacerbated in L-Dopa-treated dyskinetic mice (p < 0.05), indicating that synapse loss on D1-MSNs represents a pathological hallmark of LID (Fig. 2A). Given that microglia have been implicated as active contributors to synapse loss in other neurodegenerative diseases, we next investigated their involvement in synapse loss in PD and LID. Although IBA1 labels both microglia and macrophages within the CNS, previous work indicates that macrophages constitute less than 10% of IBA1+ cells^23^. Thus, for simplicity, we hereafter refer to IBA1+ cells as microglia, acknowledging that a minor contribution from macrophages cannot be excluded.

**Figure 2:**
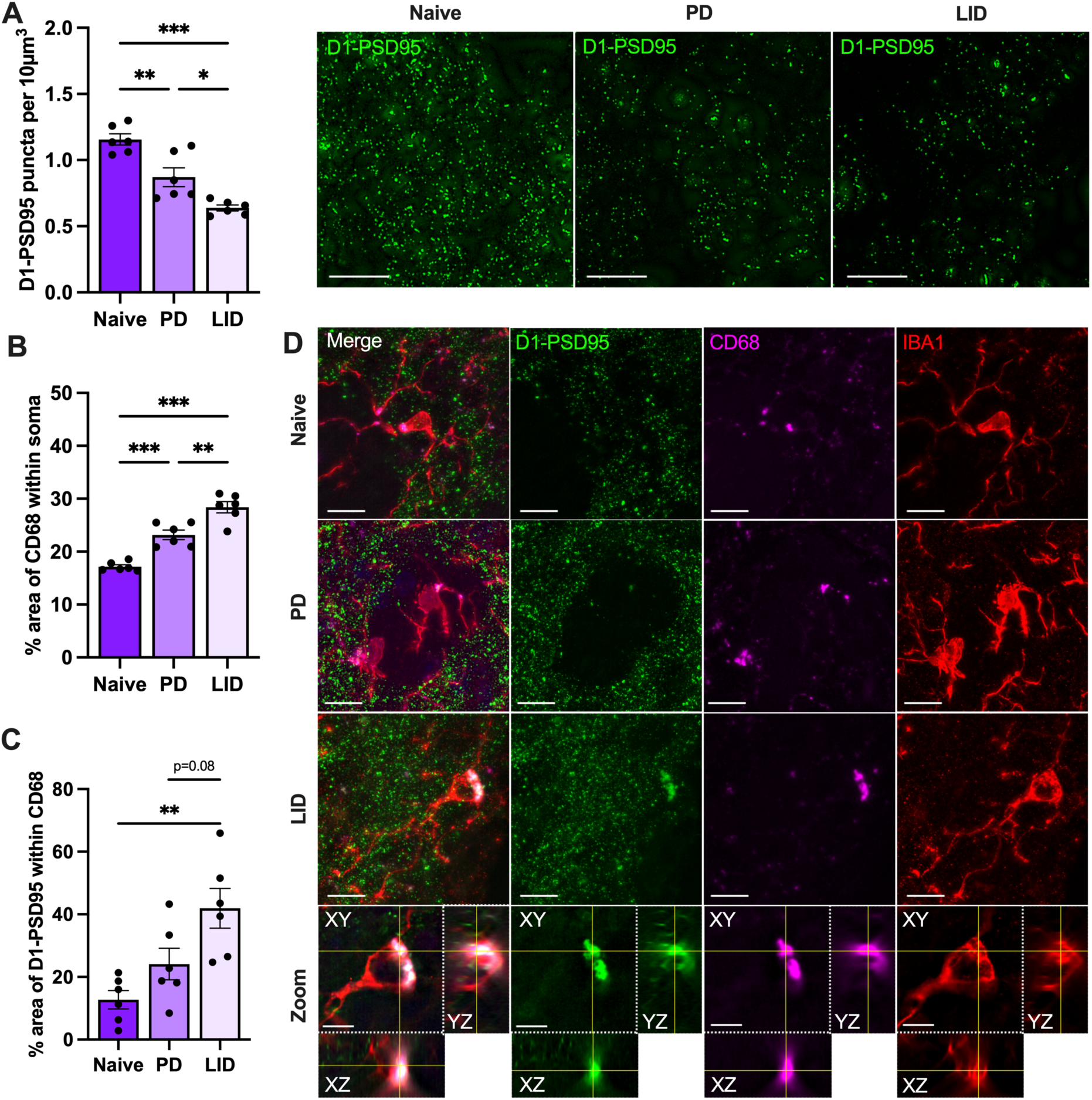
D1-PSD95-EGFP puncta loss in PD that worsens in LID is associated with microglial lysosomal accumulation of D1-PSD95 material. (A) Representative images and quantification of PSD95-EGFP puncta using SynQuant plugin in the contralateral striatum of D1-PSD95-EGFP mice showed a significant reduction in PD animals compared to naïve controls, and this loss was further exacerbated in LID mice (One-way ANOVA: F_(2,15)_=27.3, p<0.001 with Bonferroni post-hoc). (B) CD68+ lysosomal area per microglial cell body was significantly greater in PD relative to naive mice and LID mice displayed an even greater area compared to PD mice (One-way ANOVA: F_(2,15)_=46.54, p<0.001 with Bonferroni post-hoc). (C) Area of PSD95-EGFP material within CD68+ lysosomal compartments was increased in LID mice compared to naïve and PD mice (One-way ANOVA: F_(2,15)_=8.703, p=0.0031 with Bonferroni post-hoc). (D) Representative images show colocalization of D1-PSD95-EGFP synaptic material with CD68+ lysosomes of IBA1+ microglia and orthogonal views specifically show engulfment of PSD95+ inclusions within lysosomal compartments in LID mice. N = 6 per group. Scale bar = 10μm and scale bar = 5μm for zoomed in images. All values represented mean ± SEM. *p<0.05, **p<0.01, ***p<0.001.

In line with growing evidence that neuroinflammation contributed to LID^20,23^, in the current study, stereological analysis confirmed that dyskinetic mice displayed significantly higher numbers of striatal IBA1+ microglia (p < 0.05; Supplementary Fig. 1C). While the number of microglia co-expressing CD68 was not significantly different in LID mice compared to PD mice (p = 0.08; Supplementary Fig. 1D), CD68+ lysosomal area per microglial cell body was significantly greater in D1-PSD95-EGFP dyskinetic mice relative to parkinsonian mice (p < 0.01; Fig. 2B), supporting enhanced phagocytic activity as a defining feature of LID. Notably, LID mice exhibited a 3.3-fold statistically significant increase in PSD95-EGFP within the CD68+ lysosomal compartment compared to naïve mice (p < 0.01) and a 1.75-fold increase compared to PD mice (p = 0.08), indicating the presence of PSD95-EGFP material within microglial lysosomes and raises the possibility that microglial phagocytosis contributes to the observed synapse loss (Fig. 2C-D).

### Microglia-Mediated PSD95 synapse loss on D2-MSNs occurs in PD without further worsening in LID

In the striatum of D2-PSD95-EGFP mice, compared to naïve mice, we observed a trend toward significantly reduced PSD95-EGFP puncta in parkinsonian animals (p = 0.054) and a statistically significant decrease in dyskinetic mice (p < 0.01). There was no statistically significant difference between dyskinetic and parkinsonian mice (p = 0.76), indicating that synapse loss in D2-MSNs is a pathological feature of PD that does not change in LID (Fig. 3A). As observed for D1 synapses, we assessed whether microglia also contain D2-PSD95 material. Microglia of dyskinetic D2-PSD95-EGFP mice exhibited a significantly greater CD68+ lysosomal area (p < 0.001; Fig. 3B) and a significantly increased accumulation of PSD95 puncta (p < 0.05; Fig. 3C) when compared to naïve mice, supporting microglial contribution to D2 synapse loss.

**Figure 3:**
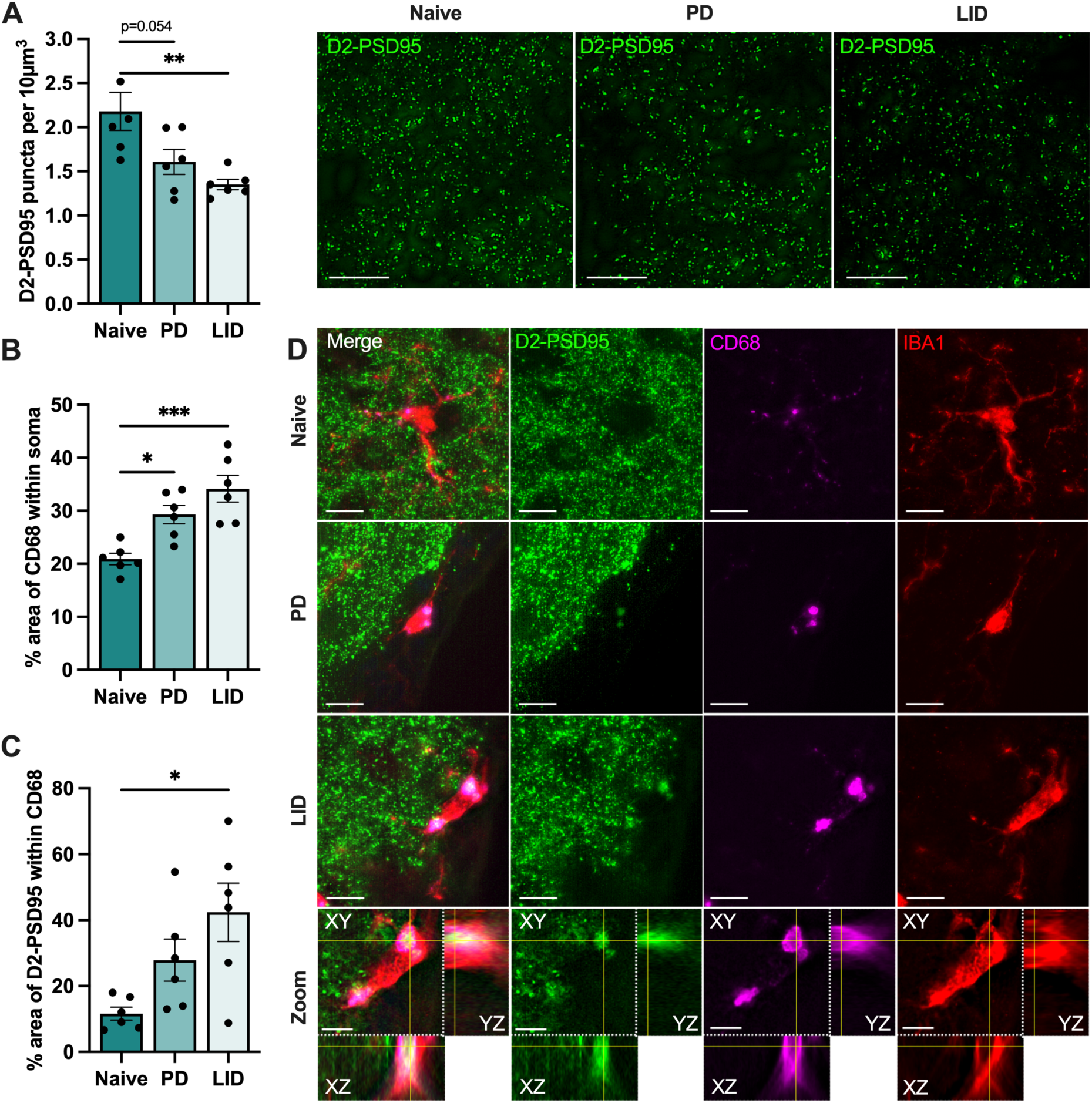
D2-PSD95-EGFP puncta loss in PD and LID is associated with microglial lysosomal accumulation of D2-PSD95 material. (A) Representative images and quantification of PSD95-EGFP puncta using SynQuant plugin in the contralateral striatum of D2-PSD95-EGFP mice showed a trend toward significant reduction in PD animals compared to naïve controls (p=0.054) and a significant decrease in LID mice (One-way ANOVA: F_(2,15)_=7.723, p=0.0049 with Bonferroni post-hoc). (B) CD68+ lysosomal area per microglial cell body was significantly greater in LID mice compared to naïve and PD mice (One-way ANOVA: F_(2,15)_=12.84, p=0.0006 with Bonferroni post-hoc). (C) Area of PSD95-EGFP material within CD68+ lysosomal compartments was significantly increased in LID mice compared to naïve mice (One-way ANOVA: F_(2,15)_=5.782, p=0.014 with Bonferroni post-hoc). (D) Representative images showed colocalization of D2-PSD95-EGFP synaptic material with CD68+ lysosomes of IBA1+ microglia, and orthogonal views specifically showed engulfment of PSD95+ inclusions within lysosomal compartments in LID mice. N = 6 per group. Scale bar = 10 μm and scale bar = 5 μm for zoomed-in images. All values represented mean ± SEM. *p<0.05, **p<0.01, ***p<0.001.

### Microglia elimination prevents synapse loss and reduces LID

Having established that LID is associated with exacerbated D1-MSN specific synapse loss, increased microglial reactivity, and accumulation of synaptic material within lysosomal compartments, indicative of phagocytic activity, we next investigated whether inhibiting microglia-mediated D1-MSN synapse removal could reduce LID severity. Accordingly, we administered the CSF1R inhibitor PLX3397 via chow to deplete microglia after the 6-OHDA lesion was fully established (confirmed lesion shown in supplementary Fig. 2A), but one week before initiating L-Dopa treatment to allow for the depletion to occur, and assessed the development of dyskinetic behaviour in microglia-depleted D1-PSD95-EGFP mice (Fig. 4A). Analysis revealed a statistically significant main effect of time (p < 0.001) and treatment (p < 0.01), with no interaction (p = 0.22), indicating PLX3397 consistently reduced LID severity while both groups exhibited progressive worsening over time (Fig. 4B). Bonferroni post-hoc tests confirmed lower AIMs scores in PLX3397-treated mice on Day 1 (p < 0.01), Day 4 (p < 0.01), Day 7 (p < 0.001), Day 10 (p = 0.06), and Day 13 (p < 0.05). We next examined whether PLX3397-induced depletion of IBA1+ microglia (88% reduction; p < 0.001), which in turn resulted in an 81% reduction of CD68+ phagocytic microglia (p < 0.001), led to preserving striatal synapses (Fig. 4C). Indeed, PLX3397-treated D1-PSD95-EGFP mice exhibited significantly more D1-PSD95 puncta (p < 0.01; Fig. 4D) compared to normal chow controls, indicating that in the absence of microglia, D1-MSN specific synapse loss induced by L-Dopa treatment is mitigated. Collectively, these findings suggest that preventing microglia-driven synapse loss on D1-MSN represents a viable pharmacological strategy to reduce LID severity in mice.

**Figure 4:**
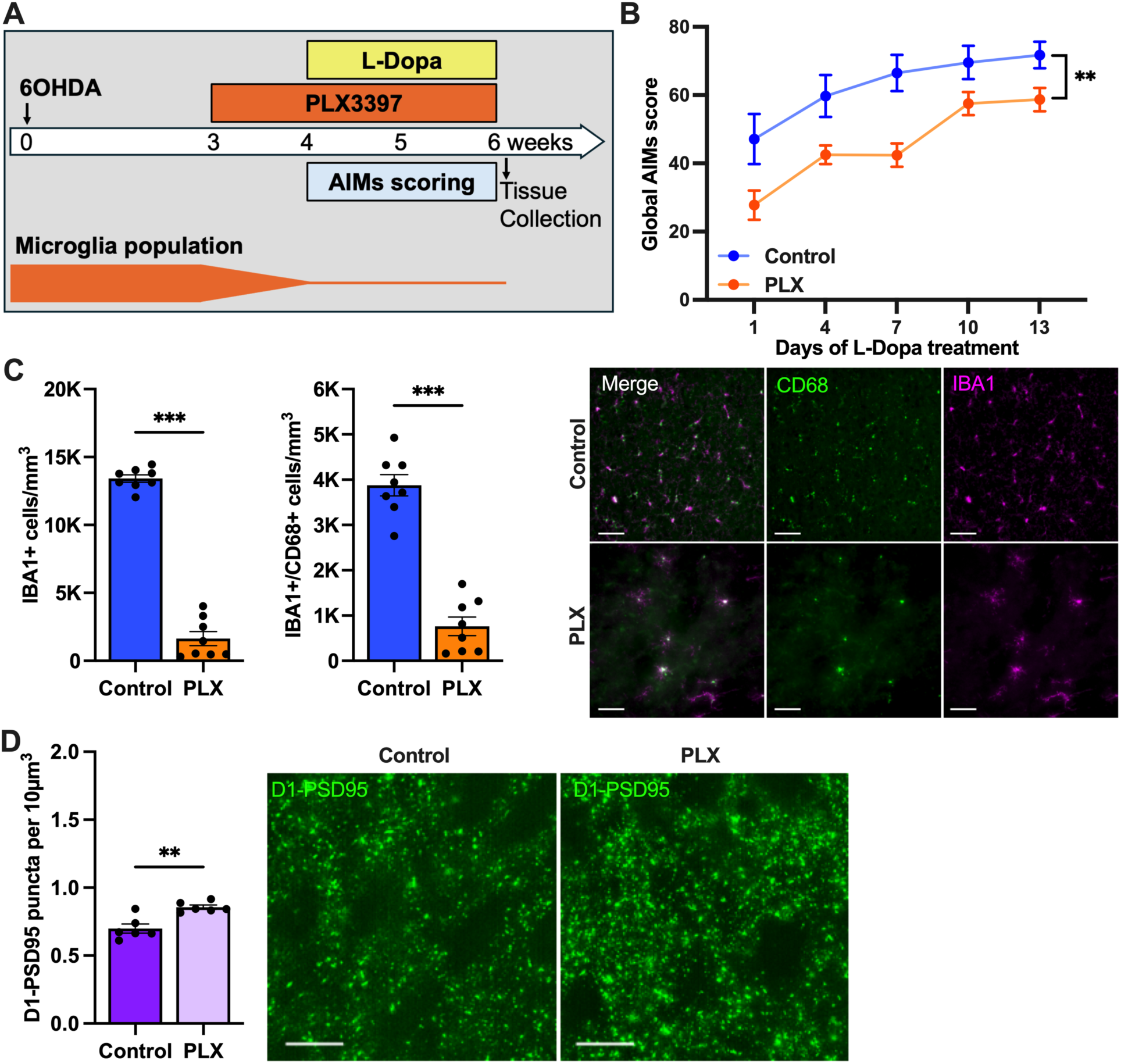
Microglia elimination prevented synapse loss and reduced LID severity. (A) Experimental timeline showing administration of PLX3397 via chow in 6-OHDA lesioned animals and AIMs scoring days. (B) Behavioural testing revealed PLX3397 treatment significantly reduced AIMs scores (Two-way repeated measures ANOVA: Interaction F_(4,124)_=1.428, p=0.2287; Time F_(4,124)_=29.67, p<0.0001; Treatment F_(1,31)_=10.03, p=0.0034 with Bonferroni post-hoc tests). (C) Stereological analysis revealed PLX3397 treatment significantly reduced IBA1+ microglia (Unpaired t-test: t=20.2, df=14, p<0.001) and CD68+ phagocytic microglia compared to controls (Unpaired t-test: t=9.98, df=14, p<0.001). (D) PLX3397-treated D1-PSD95-EGFP mice exhibited significantly more D1-PSD95 puncta compared to controls (Unpaired t-test: t=4.294, df=10, p=0.0016). N=16 for control and N=17 for PLX3397 for behavioural analysis; N=8 per group for microglia quantification; N=6 per group for synapse analysis. Scale bar = 50 μm for microglia images and scale bar = 10 μm for synaptic puncta images. All values represent mean ± SEM. *p<0.05, **p<0.01, ***p<0.001.

### Microglial turnover through elimination and repopulation prevents synapse loss and reduces LID

Microglia rapidly repopulate following discontinuation of PLX3397 treatment, and this process has generated considerable interest as a potential therapeutic strategy for CNS disorders, including AD and PD. We first established the timeframe required for microglia to return to baseline levels after depletion. Consistent with previous reports, by one week post PLX3397 removal, microglial numbers exceeded those of control mice by approximately 62%, and these cells exhibited a more reactive phenotype (Fig. 5A). By five weeks, microglial numbers had fully stabilized, consistently matching pre-depletion levels both in number and morphology (Fig. 5A). Accordingly, we selected a five-week repopulation period for subsequent experiments.

**Figure 5:**
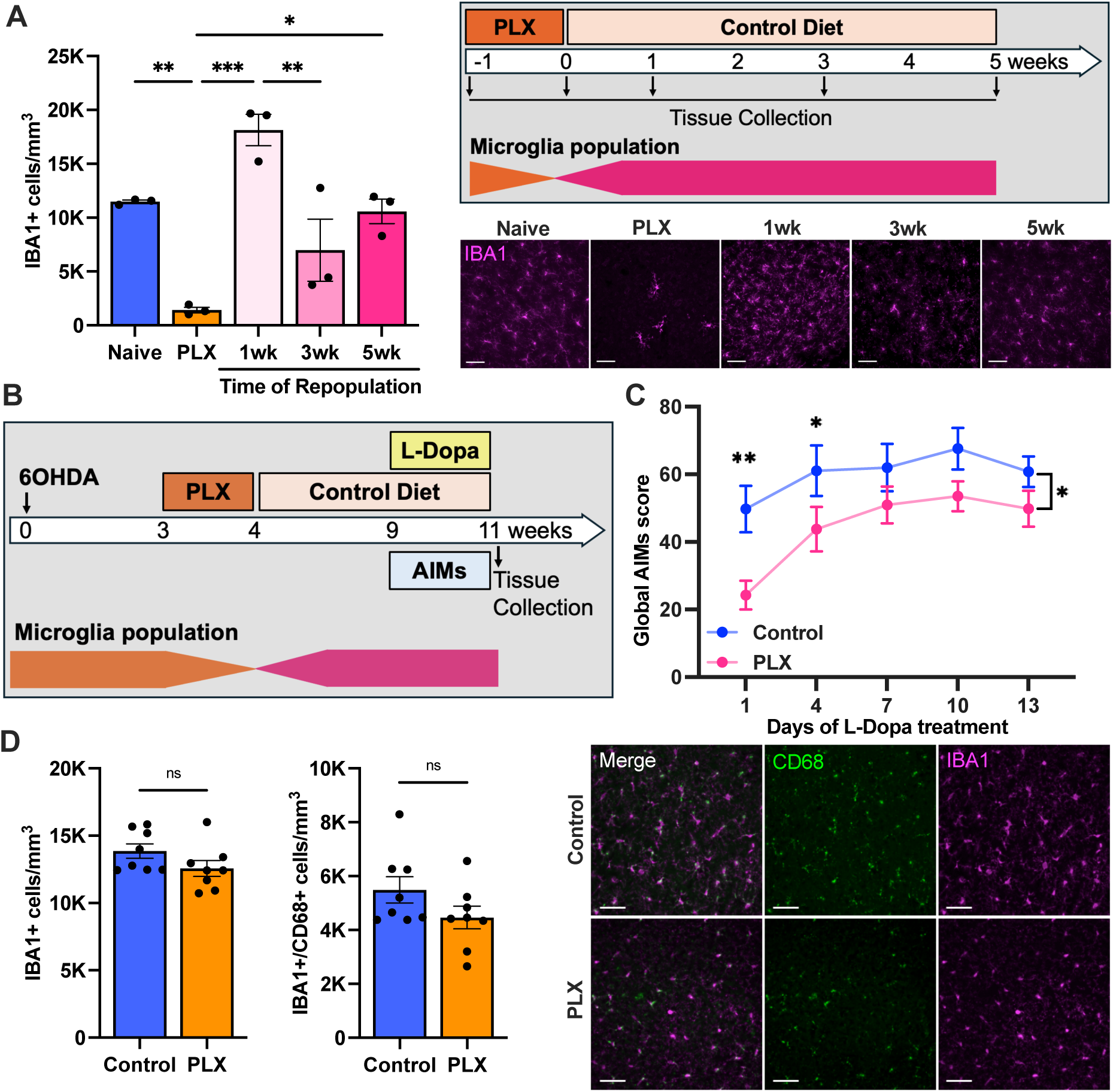
Microglia repopulation following PLX3397 withdrawal reduced LID severity. (A) Experimental timeline, representative images, and stereological quantification revealed time course of microglia repopulation in the striatum of C57BL/6J mice after withdrawal of PLX3397, with microglial numbers exceeding those of control mice by 1 week, and by 5 weeks, numbers consistently returned to baseline levels (One-way ANOVA: F_(4,10)_=15.90, p=0.0002 with Bonferroni post-hoc analysis). (B) Experimental timeline to investigate the anti-dyskinetic efficacy of PLX3397 depleted and repopulated microglia in 6-OHDA lesioned mice. (C) Behavioural scoring revealed PLX3397 repopulated microglia significantly reduced AIMs scores (Two-way repeated measures ANOVA: Interaction F_(4,112)_=1.904, p=0.1147; Time F_(4,112)_=17.38, p<0.0001; Treatment F_(1,28)_=4.560, p=0.0416 with Bonferroni post-hoc tests). (D) Representative images and quantification of IBA1+ and CD68+ phagocytotic microglia after repopulation showed no significant differences between PLX3397-treated and control mice (IBA1: Mann-Whitney U=16, p=0.10; CD68: Mann-Whitney U=20, p=0.22). N=3 for repopulation time course; N=15 per group for behavioural analysis; N=8 per group for microglia quantification. Scale bar = 50 μm. All values represent mean ± SEM. *p<0.05, **p<0.01, ***p<0.001.

We hypothesized that replacing reactive microglia associated with the 6-OHDA lesion (lesion severity confirmed; Supplementary Fig. 2B) with newly repopulated, non-reactive microglia prior to initiating L-Dopa treatment would mitigate microglia-mediated synapse loss and reduce the development of LID (Fig. 5B). Indeed, analysis revealed a significant main effect of time (p < 0.0001) and treatment (p < 0.05), with no interaction (p = 0.11), indicating that PLX3397 treatment reduced LID severity, compared to animals that received control diet throughout, while both groups developed more AIMs over time (Fig. 5C). Bonferroni post hoc tests confirmed lower AIM scores in PLX3397-treated mice on Day 1 (p < 0.01), Day 4 (p < 0.05), and a trend toward significance on Day 10 (p = 0.09).

We next aimed to confirm that these renewed microglia indeed had a less reactive phenotype. However, we did not find a statistically significant difference between control and PLX3397-treated mice in either total striatal IBA1+ or CD68+ phagocytotic microglia numbers (Fig. 5D). This may be because tissue was collected after 14 days of L-Dopa treatment, at which point L-Dopa had once again led to microglial activation. Lastly, we assessed whether the observed behavioural benefit of repopulated microglia was associated with reduced synapse loss. In the striatum of D1-PSD95-EGFP mice, we observed significantly more PSD95-EGFP puncta (p < 0.05; Fig. 6A) in mice with repopulated microglia compared to controls. Furthermore, these repopulated microglia exhibited a significantly reduced CD68+ lysosomal area (p < 0.05; Fig. 6B) and a significantly lower amount of PSD95 puncta within microglia (p < 0.05; Fig. 6C-D) relative to control mice. These findings suggest that repopulated microglia are initially less phagocytic and that preserved D1-PSD95 synapses coincide with less severe LID.

**Figure 6:**
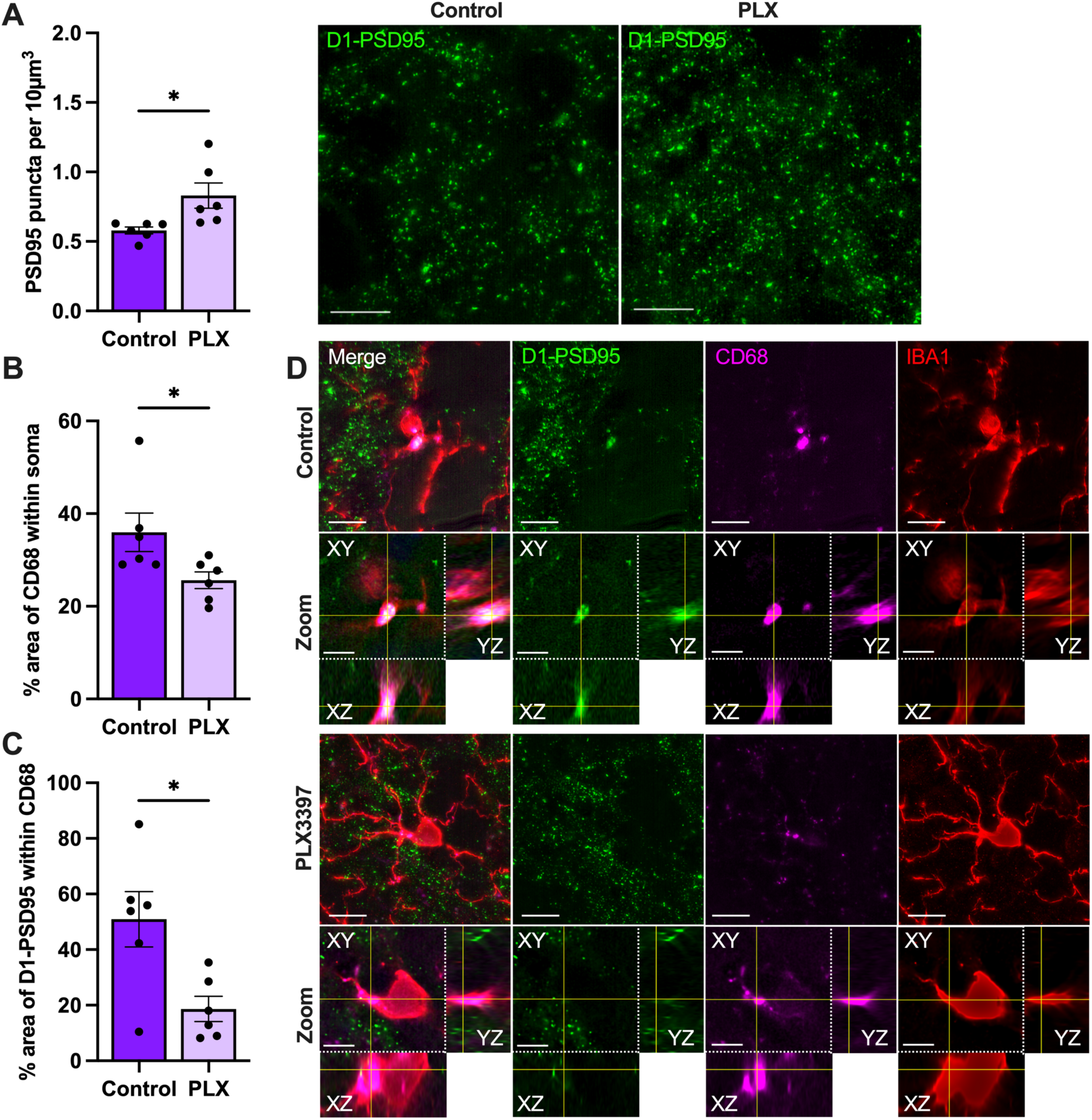
Repopulated microglia preserved D1-PSD95 synapses and exhibited reduced phagocytic activity. (A) Representative images and quantification of PSD95-EGFP puncta using SynQuant in the striatum of D1-PSD95-EGFP mice showed significantly more puncta in mice with repopulated microglia compared to controls (Unpaired t-test: t=2.65, df=10, p=0.02). (A) CD68+ lysosomal area per microglial cell body was significantly reduced in repopulated microglia compared to controls (Mann-Whitney U=3, p=0.02) and (C) area of PSD95-EGFP material within CD68+ lysosomal compartments was significantly lower in repopulated microglia compared to controls (Unpaired t-test: t=2.95, df=10, p=0.01). (D) Representative images show colocalization of D1-PSD95-EGFP puncta with CD68+ lysosomes in control mice and reduced lysosomal PSD95 inclusions in mice with repopulated microglia and orthogonal views specifically show engulfment of PSD95+ inclusions within lysosomal compartments in control mice that is not present in mice with repopulated microglia. N=6 per group. Scale bar = 10 μm; zoomed-in images = 5 μm. All values represent mean ± SEM. *p<0.05

## Discussion

Microglia mediated synapse loss underlies multiple neurodegenerative diseases, and therapeutic strategies to prevent this process present a major focus of research. Since the striatum, a region affected in PD and LID, contains two intermingled MSN subtypes with opposing roles, distinguishing between them is critical. To address this, we generated novel D1-PSD95-EGFP and D2-PSD95-EGFP reporter strains to enable MSN-specific synapse visualization. In the current study, we report four major findings. First, both D1- and D2-MSNs exhibited significant loss of PSD95-containing synapses in PD, with synapse loss in D1-MSNs further exacerbated in LID. Second, this synapse loss coincided with markedly increased PSD95 material within lysosomal compartments of microglia in LID, implicating active phagocytosis. Third, depleting microglia using the CSF1R inhibitor PLX3397 mitigated D1-MSN synapse loss and reduced LID severity. Lastly, when a microglia depletion - repopulation paradigm was applied that replaced reactive microglia with newly repopulated cells, LID severity was attenuated, D1-MSN synapses were preserved, and synaptic material within microglia was reduced. Collectively, these findings demonstrate that striatal synapse loss in D1-MSNs is a critical contributor to LID pathogenesis and that preventing microglia-mediated synapse removal represents a promising therapeutic strategy.

### PSD95 synapses on D2-MSNs are lost in PD and not restored in LID

Previous studies have reported that D2-MSN spines lost in the PD state are restored during LID ^5–9^, however, in the current study, we observed that D2-PSD95 synapse loss occurs in PD and remains unchanged in LID. A recent investigation provides a compelling explanation for these discrepancies^10^. First, our results can be explained as they demonstrated that tissue collection timing is critical. Spine regrowth appears only when tissue is sampled 24 - 48 hours after the last L-Dopa dose, as in earlier reports, whereas tissue collection during peak LID expression, as in our study, reveals no difference between PD and LID^10^. More importantly, follow-up experiments in previous studies suggest that apparent spine regrowth reflects methodological limitations rather than true structural remodelling. Indeed, changes in spine density across on/off states likely result from variations in spine size, in line with a previous study showing changes in PSD95 length in LID^6^. As such, larger spines are more easily detected and smaller spines fall below the resolution threshold of traditionally used 2PLSM. This hypothesis was confirmed using a viral labelling approach combined with confocal imaging, showing that previously reported spine regrowth was largely diminished^10^. This supports the view that what was once thought to be spine regrowth on D2-MSNs may instead be synaptic strength modulation, resulting in changes in spine size rather than the formation of new spines. Our findings further confirm this interpretation, as our high-resolution PSD95 imaging indicates that D2-MSN synapse loss persists in LID, aligning with the notion that structural recovery does not occur.

Furthermore, we employed a fundamentally different approach using high-resolution PSD95 imaging to directly quantify synaptic puncta rather than relying on spine morphology. This method offers several advantages: (i) PSD95 puncta provide a functional proxy for excitatory synapses, reducing ambiguity associated with varying spine morphology; (ii) imaging can be performed with high-resolution microscopes, minimizing detection bias for small synapses and (iii) quantification of PSD95 puncta using SynQuant allows for minimum and maximum PSD95 area selection and since PSD95 area correlates with synaptic strength^34,35^, this provides a practical proxy for assessing synaptic strength without electrophysiological recordings.

### PSD95 synapse loss on D1-MSNs occurs in PD and is exacerbated in LID

While previous studies have attributed spine loss in D1-MSNs either exclusively to PD or to LID, our findings support both interpretations. Specifically, we observed a significant reduction in D1-PSD95 puncta in PD, which was further exacerbated in LID, indicating that synapse loss contributes to both PD and LID pathology. Discrepancies among earlier reports may reflect methodological differences, including species (mouse^9^ vs rat^36^), 6-OHDA injection sites (MFB^7^ vs striatum^5,6^), and lesion development timeframes (3–4 weeks vs 8 weeks^11^), all of which influence lesion severity and disease progression. Notably, prior studies consistently quantified dendritic spines using BAC transgenic mice (BAC-Drd1a or BAC-Drd2) combined with dye-filling and fluorescent imaging, focusing on individual dendritic segments. In contrast, our approach employed high-resolution imaging to quantify PSD95 puncta across striatal regions, enabling assessment of global synaptic density rather than neuron-specific spine counts. This distinction is critical, as reduced synaptic density in our study can arise from multiple structural changes such as loss of synapses per dendrite, a reduction in dendritic length, or even neuronal loss. Each of these factors would ultimately contribute to a global reduction in striatal synaptic density. Although neuronal loss of D1-MSNs has not been reported^37^, dendritic lengths are indeed reduced in PD and LID^7^, suggesting that cumulative dendritic and spine loss could amplify overall striatal synapse loss. In support of this, Fieblinger et al. demonstrated that when accounting for reduced dendritic length, total spine loss occurs in PD and worsens in LID, consistent with our findings^7^. Collectively, these observations indicate that synapse loss on D1-MSNs is a defining feature of both PD and LID, underscoring the need for therapeutic strategies aimed at preserving striatal synaptic integrity.

*Microglia phagocytose synapses lost from D1- and D2-MSN in lysosomal compartments* Synapse loss is a hallmark of multiple neurodegenerative diseases (reviewed in ^38^), and microglia-mediated synaptic phagocytosis is increasingly recognized as a key mechanism underlying this pathology. Conditions in which microglia facilitate synapse loss are consistently associated with microgliosis^38^, and PD^16–18^ and LID^19–23^ similarly exhibit pronounced microglial activation, highlighting the relevance of investigating microglia-mediated synapse removal in these disorders. In the current study, we confirmed that total and reactive microglia numbers are significantly increased in LID. Furthermore, lysosomal area per microglial cell was enlarged in PD and further exacerbated in LID, indicating enhanced phagocytic activity during dyskinesia. Interestingly, we demonstrated that these lysosomal compartments contain D1- and D2-PSD95 material, and orthogonal views confirmed true colocalization within CD68-positive lysosomes rather than an artifact of maximum projection imaging. These data suggest that the synapse loss we reported may have been facilitated by microglia. Recent reports have raised concerns that lipofuscin^39,40^, aggregates of oxidized lipids, misfolded proteins, and metals that autofluoresce across the spectrum could be misinterpreted as synaptic material. However, we are confident that our observations reflect PSD95 inclusions for two reasons. First, lipofuscin accumulation does not increase with microglial activation^40^ and would therefore be expected to remain constant across groups. Second, Stillman et al. reported that lipofuscin occupies only 1–2% of microglial area at this age, whereas our quantification revealed PSD95 material comprising ∼11% of IBA1-positive cell area in LID, far exceeding expected lipofuscin levels^40^. Collectively, these findings strongly support that microglia contain increased amounts of synaptic material in PD and LID, correlating with the synapse loss observed in our study.

### Microglia elimination prevents D1-MSN synapse loss and LID

Microglial driven excessive synaptic pruning can lead to behavioural impairments in HD^15^ and AD^12^, suggesting that preventing this process may offer a therapeutic strategy. One approach that has gained attention is blocking CSF1R signaling^41,42^, which eliminates microglia and thus abolishes activated microglia which are associated with neurodegenerative diseases. In the current study, microglial depletion using PLX3397 prevented both synapse loss and LID. This effect aligns with findings in the 5xFAD mouse model of AD^43^, where microglial elimination using PLX5622 mitigated synaptic protein downregulation and improved cognition in the Morris Water Maze. Similarly, in PD models, microglial depletion using PLX3397 reduced neurodegeneration and improved motor performance in the forced swim test^44^. Collectively, these observations, together with our findings, suggest that blocking neurodegeneration and synapse loss in particular, by eliminating the cell facilitating it, can prevent the development of LID.

### Microglia repopulation as a feasible therapeutic to prevent synapse loss and LID

Although microglia lose their homeostatic molecular signature and become activated in neurodegenerative diseases, in the healthy brain microglia also perform essential functions such as clearance of extracellular debris and maintenance of synaptic health. Therefore, prolonged microglial elimination in humans as a therapeutic approach may be questionable. Interestingly, microglia can rapidly repopulate, and previous studies suggest that repopulated microglia may reset pathological states by replacing them with functionally intact cells^25–27^. In the current study, mice harbouring repopulated microglia developed less severe LID, and this behavioural improvement was associated with the prevention of synapse loss and reduced synaptic material within microglia. One caveat was that although CD68+ area per microglial cell decreased, consistent with a less reactive phenotype, the total number of IBA1+ and CD68+ microglia in the striatum was not significantly reduced. We propose that 14 days of L-Dopa treatment likely reactivated microglia over time. This interpretation aligns with findings in the 5xFAD model of AD, where microglial repopulation in the short term resulted in less reactive cells, improved cognition in the Morris Water Maze, reduced PSD95 puncta in the hippocampus, and decreased synaptic inclusions within microglia^45^. However, prolonged exposure to the AD environment in that study led repopulated microglia to revert to a pathological state, suggesting that the beneficial effect may not be sustained indefinitely. Consequently, therapeutic benefits to prevent LID may require repeated cycles of depletion and repopulation. Collectively, microglial repopulation offers a promising strategy to prevent microglia-mediated synapse loss and improve behavioural outcomes in neurodegenerative diseases. Given the proven efficacy of PLX3397 in human studies^27^, microglia repopulation paradigms in parkinsonian patients on L-Dopa appears feasible. Our findings provide translational potential and a mechanistic basis for future clinical investigations.

## Conclusion

In conclusion, this study introduces novel D1-PSD95-EGFP and D2-PSD95-EGFP reporter mouse lines, enabling high-resolution visualization of MSN subtype-specific synapses and providing a powerful platform to interrogate striatal connectivity in PD and LID. Using these models, we demonstrate that synapse loss occurs in both MSN subtypes in PD, with additional exacerbation on D1-MSNs during dyskinesia, and that this loss coincides with microglial engulfment of synaptic material. Furthermore, by demonstrating that both depletion and subsequent repopulation of microglia can attenuate dyskinetic behaviours and preserve striatal synaptic integrity, this study provides mechanistic insight into how resetting microglial states may counteract maladaptive synaptic remodelling. Collectively, these observations offer a compelling rationale for translating microglial modulation into clinical interventions aimed at mitigating motor complications in LID.

## Notes

### Competing Interest Statement

The authors have declared no competing interest.

### Funding

- Australian Government Research Training Program Scholarship
- The Helen and David Baffsky Fellowship
- The Boyarsky Family
- Andrew Michael and Michele Brooks
- John and Debbie Schaffer
- Richard Gelski
- Alex Sundich and Bridge Street Capital Partners
- Doug Battersby and family
- David King and family
- Harry Holden
- Tony and Vivian Howland-Rose
- The ISG Foundation
- Stanley and Charmaine Roth
- Richard, Adrian and Tom O'Connor
- Marnie and Gary Perlstein
- David Schwartz
- Stephen Young
- Nelson Meers Foundation
- McCusker Foundation

