## Supplementary Figure 1 for "Novel PSD95 reporter mice reveal medium spiny neuron subtype-specific synapse loss in PD and L-dopa induced dyskinesia and identify microglia mediated synapse removal as a therapeutic target for dyskinesia"

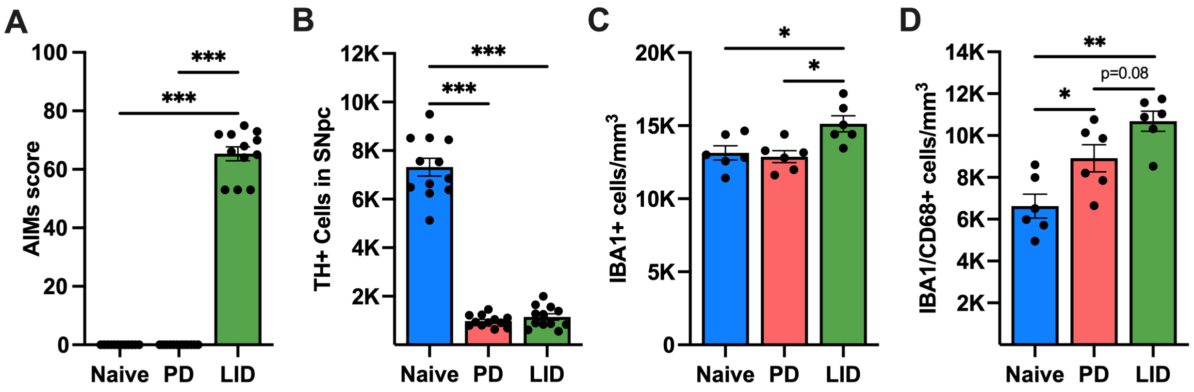


**Supplementary Figure 1: Disease stage-dependent LID development, 6-OHDA lesion severity and microglial quantification.** (A) Global AIMs scores confirmed that only mice receiving L-Dopa and obtaining an AIMs score above 40 were categorized as LID and included for analysis (One-way ANOVA: $F_{(2,33)}$=773.2, p<0.001 with Bonferroni post-hoc analysis). (B) Stereological quantification of TH+ neurons in the SNpc confirmed extensive DA neuron loss in PD and LID groups compared to naïve controls (One-way ANOVA: $F_{(2,33)}$=257.6, p<0.001 with Bonferroni post-hoc analysis). (C) Stereological analysis showed significantly higher numbers of striatal IBA1+ microglia in dyskinetic mice compared to naïve controls and PD mice (One-way ANOVA: $F_{(2,15)}$=6.324, p=0.0102 with Bonferroni post-hoc analysis) and (D) CD68+ reactive microglia were significantly increased in PD mice compared to naïve controls with a trend towards a significant further increase between PD and LID (One-way ANOVA: $F_{(2,15)}$=12.74, p=0.0006 with Bonferroni post-hoc analysis). N=6 per group for microglial analysis; N=12 per group for behavioural and TH quantification. All values represent mean ± SEM. *p<0.05, ***p<0.001.
