## Supplementary Figure 2 for "Novel PSD95 reporter mice reveal medium spiny neuron subtype-specific synapse loss in PD and L-dopa induced dyskinesia and identify microglia mediated synapse removal as a therapeutic target for dyskinesia"

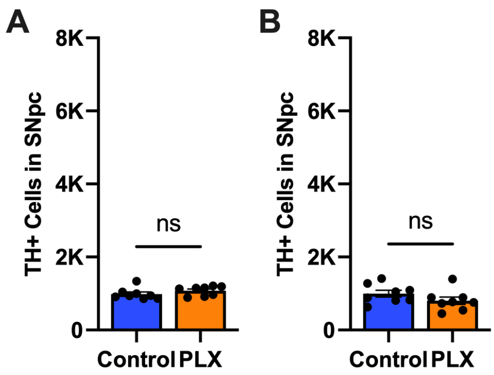


**Supplementary Figure 2: Consistent 6-OHDA lesion severity in PLX3397 microglia depleted and repopulated mice.** Extensive dopaminergic lesions were confirmed, as there were no statistically significant differences between control and (A) PLX3397 microglia depleted (Unpaired t-test: t=1.27, df=14, p=0.22) and (B) PLX3397 treated and repopulated microglia mice (Unpaired t-test: t=1.42, df=14, p=0.18). N=8 per group. All values represent mean ± SEM.
